# Spontaneous intake and long-term effects of essential oils after a negative postnatal experience in chicks

**DOI:** 10.1101/452136

**Authors:** Laurence A. Guilloteau, Anne Collin, Alexia Koch, Christine Leterrier

**Affiliations:** BOA, INRA, Université de Tours, 37380 Nouzilly, France; PRC, CNRS, IFCE, INRA, Université de Tours, 37380 Nouzilly, France

**Author notes:** Correspondence: Dr. Laurence Guilloteau.

**Keywords:** Essential oil, self-medication, broiler, chicks, postnatal experience

## Abstract

The postnatal period is critical for broiler chicks as they are exposed to, possibly stressful, environmental changes in the hatchery and during transportation to the rearing houses. The ability of broiler chicks to spontaneously drink essential oils (EO) to mitigate the effects of a negative postnatal experience was tested. Chicks were either immediately placed in the rearing facility (C group), or subjected to a 24h-delay period before their placement (D group), mimicking the possible transportation delay in commercial conditions.

In experiment 1, each group had access to either water only or to water and one EO (cardamom, marjoram or verbena) from D1 to D13. The verbena EO intake was higher in the D group than in the C group from D1 to D6 and the cardamom EO intake was lower in the D group than in the C group from D6 to D13.

In experiment 2, half of the groups had access to water only and the other half was offered water and the 3 EO simultaneously. The EO were not differently chosen by chicks between D and C groups except a lower cardamom EO intake was observed in the D group than in the C group from D6 to D12. The delayed placement of the D group reduced chicken growth until 34 days of age and temporarily increased the feed conversion ratio, but did not affect their welfare or the prevalence of health disorders. The EO intake did not allow the chicks in the D group to overcome the growth reduction, but did overcome the reduction in *Pectoralis major* muscle yield. In conclusion, chicks are able to make spontaneous choices regarding EO intake according to their postnatal experience when EO are presented individually, but in our experimental design, they were not when EO were simultaneously presented. The EO intake only partially mitigated the decrease in chicken performance after the negative postnatal experience.

## Introduction

The postnatal period is a critical period for livestock animals. They have to cope with specific management conditions, and exposure to adverse environmental conditions that may result in stress responses. Stress during early life can induce persistent changes in physiology, behavior, and immune phenotype (Pryce et al., 2002). Strengthening the robustness ‐that is to say the capacity of the animal to adapt to environmental disturbances‐ during the postnatal period is a potential strategy to reduce the immediate and long-lasting effects of stressful early experiences. It can also contribute to the improvement of the animal’s sanitary status and to the reduction in the use of antimicrobial drugs. One approach originally observed in wild animals is the stimulation of self-medication behavior (SM) or zoopharmacognosy. It has been defined as the ability of animals to select and use specific plants or substrates with medicinal properties to control or to prevent diseases (Rodriguez and Wrangham, 1993) or situations of discomfort. Another view of self-medication is defined by Forbey et al (Forbey et al., 2009) as a homeostatic behavior. In farm animals, observations of SM were reported in ruminants (Grade et al., 2009) and research was mainly focused on plants associated with anti-parasitic properties (Villalba et al., 2014;Ventura-Cordero, 2018).

The objective to reduce the use of chemical antimicrobial drugs in farm animals encourages the proposal of alternatives to these medicines (Murphy, 2017). Essential oils (EO) extracted from medicinal plants have multi-functional properties including antimicrobial, antioxidant, immunostimulatory, anti-inflammatory, and nervous system regulatory properties (Bakkali et al., 2008;Adorjan and Buchbauer, 2010;Dobetsberger and Buchbauer, 2011;de Sousa et al., 2015). These properties are related to the composition of the EO, mainly terpenoids (monoterpenes, sesquiterpenes) and a variety of aromatic compounds. Phenols, alcohols, ketones, and aldehydes are mainly associated with antibacterial actions (Nazzaro F, 2013). Phenylpropanoids (de Cassia da Silveira et al., 2014) and terpenoids such as the oxide 1,8-cineole are known to have anti-inflammatory effects, and positive effects on the digestive and respiratory systems (Juergens, 2014;Adaszynska-Skwirzynska and Szczerbinska, 2017).

In chickens, EO have received attention as growth and health promoters and have been used as feed additives (Brenes and Roura, 2010;Alleman et al., 2013;Gabriel et al., 2013;Zeng et al., 2015;Adaszynska-Skwirzynska and Szczerbinska, 2017). In these studies, EO were included in feed and chickens did not have the choice to ingest them or not. If chickens are able to select EO with medicinal effects that are the most adapted to them in a challenging situation, this would potentially improve their robustness and reduce the use of drugs.

To test the hypothesis that chicks are able to spontaneously consume EO according the discomfort they have experienced, we have developed an experimental setting to reproduce the adverse conditions where they are subjected to during the postnatal period. In poultry production systems, chicks undergo transportation from the hatchery to the rearing houses and stressors like temperature variations, jolts in transportation boxes due to truck movements, feed and water deprivation lasting between several hours and 2 or 3 days after hatching. Feed and water deprivation in chicks has long-lasting effects on performance (Bigot et al., 2003;Gonzales et al., 2003;de Jong et al., 2017), and also on physiological and immune parameters (Gonzales et al., 2003;Shakeel et al., 2016), which can result in higher susceptibility to diseases and mortality (Shakeel et al., 2016). The long-lasting effects of post-hatch transportation have also been described in terms of chick behavior, health, and performance (Zulkifli et al., 1994;Valros et al., 2008;Oviedo-Rondon et al., 2009;Bergoug et al., 2013;Jacobs et al., 2016).

In this study, two experiments were performed. The first experiment was designed to assess the capacity of chicks to spontaneously ingest EO and to analyze whether this intake was related to their postnatal experience. The second experiment was designed to assess the capacity of chicks to choose between three EO in free access and in addition to the drinking water, as well as to observe the kinetics of EO choice, and to analyze the effects of the EO on the performance, welfare, and health of the chickens over the whole growing period.

## Materials and Methods

All procedures used in these experiments were approved by the local ethics committee (Comité d’Ethique en Expérimentation Animale Val de Loire, Tours, France; permission no 01730.02 and 2015070815347034v2, APAFIS#1082) and carried out in accordance with current European legislation (EU Directive 2010/63/EU).

### Model of a postnatal negative experience in chicks

After hatching, chick transportation to the broiler farms can occur in suboptimal conditions (Zulkifli et al., 1994;Valros et al., 2008;Oviedo-Rondon et al., 2009;Bergoug et al., 2013;Jacobs et al., 2016). To analyze the consequences of this experience over the whole growing period, eggs (Hubbard Classic^®^, Quintin, France) were incubated for 21 days under standard conditions. After opening the incubator (T0), the chicks were sex-sorted according their plumage, wing-tagged, and immunized with a vaccine against infectious bronchitis (IB) (NOBILIS IB 4/91^®^, Intervet, Beaucouzé, France) by the conjunctival route. Afterwards the chicks were either placed in pens in the rearing facility after their withdrawal from the incubator (Control group, C) or were submitted to a 24h-delay period before their placement (Delayed group, D). The latter were deprived of feed and water and put in transportation boxes that underwent irregular movement and variable room temperature: 32°C (30 min), 21°C (90 min), 32°C (30 min) and then at 21°C with alternating cycles of box movement (M) and immobility (I) for 24h after the C group was placed in pens. One cycle was 45 min (M), 15 min (I), 30 min (M), 30 min (I). These conditions were combined to be the closest to the actual suboptimal conditions in broiler chicken livestock. Chicks were allotted according to the time of hatching (50% that hatched in the incubator more than 12 hours before T0, and 50% that hatched between 12h and 0h (= T0) and sex (50% male/50% female as determined at T0). Chicks were reared at the Pôle d’Expérimentation Animale de Tours (PEAT) (INRA Centre Val de Loire, France) in standard temperature and light conditions with *ad libitum* access to water and with a wire mesh platform and a perch for environmental enrichment. At D13, the chickens were transferred to another livestock building and placed in larger pens (2m × 1m) for the growth phase until D34. They had *ad libitum* access to feed without anticoccidial drugs. They were fed with a standard starting diet (metabolizable energy = 12.8 MJ/kg, crude protein = 22%) until 19 days and then a rearing diet from 19 to 34 days.

### Essential oils

The essential oils (EO) were chosen for their common and complementary properties to control infectious challenges, to reduce the stress response, and to improve the functions of the digestive and immune systems. Three EO were chosen based on scientific literature, expert advice from practitioners and preliminary results from experiments performed with 12 EO in chickens. Cardamom (*Elettaria cardamomum*) (1480CQ, batch S12A, Herbes et Traditions, Comines, France), marjoram (*Origanum majorana*) CT thujanol (2507CQ, batch S12D, Herbes et Traditions), and lemon verbena (*Lippia citriodora*) (FLE094, batch H181013MA, Florihana, Caussols, France) were used to assess the spontaneous intake of EO in the C and D groups. Each EO was diluted in water (10mg/L) and delivered in a bottle. The main components obtained by gas chromatography coupled to mass spectrometry are listed in Table 1.

**Table 1.**
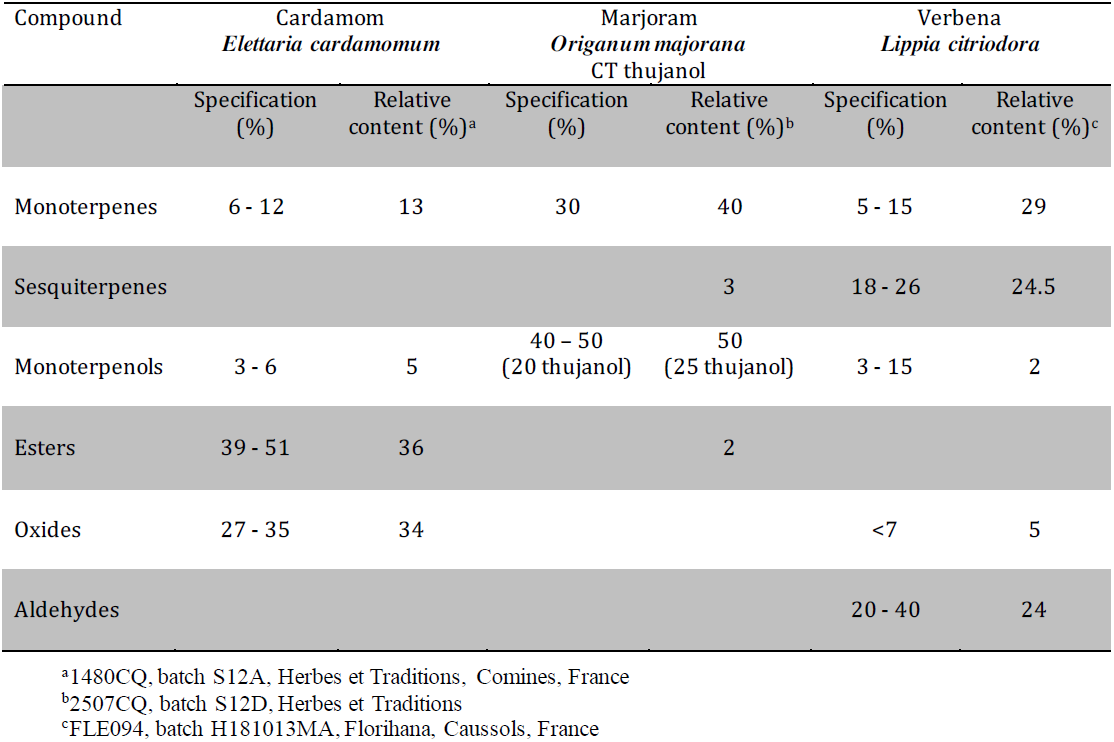
Essential oils composition

In addition to their antimicrobial and antioxidant activities (Singh et al., 2008;Choupani M, 2014;Bina and Rahimi, 2017), these EO have complementary properties. Cardamom EO has been demonstrated to have antispasmodic and anti-inflammatory activities (al-Zuhair et al., 1996), and gastroprotective properties (Jamal et al., 2006). Among various biological activities, marjoram EO has hepatoprotective activities (Bina and Rahimi, 2017). Lemon verbena EO is reputed to have analgesic, anti-inflammatory, sedative, and digestive properties (Pascual ME, 2001).

Based on previous studies (Galal et al., 2016), each EO was diluted in water (0.001%) and delivered in a bottle. The main components obtained by gas chromatography coupled to mass spectrometry in each EO are listed in Table 1.

### Experimental design

#### 1. Experiment 1

After incubator opening, 192 chicks were placed in pens (0.5m × 1m) in the C group (n = 96) or D group (n = 96). Feed and water (W) supplies were provided for chicks in free *ad libitum* access in the pens from D0 to D13 for the C group and from D1 until D13 post-hatching for the D group. Each essential oils (EO) was placed at D1 in the EO-C and EO-D groups (six pens for each EO). Chicks were allocated to the W-C or W-D, or EO-C or EO-D group (six pens/group, four chicks/pen). Two bottles, one containing water and one containing one EO were available per pen for the EO groups. Two bottles of water were available for the W groups (Figure 1). The bottle position was changed every day for a week and every 2-3 days during the second week to prevent the chicks from getting used to the position of the bottles. The intake of water and of each EO was recorded at D1, D2, D4, D6, D9 and D13. Water and EO were changed each time of intake recording and supplemented if necessary between two measures of intake. Water and EO intakes were first compared between groups for the 13 days post hatching when EO were provided. Water and EO intakes have been then expressed as a percentage of the total liquid intake since differences in total liquid intakes were observed between C and D groups.

**Figure 1.**
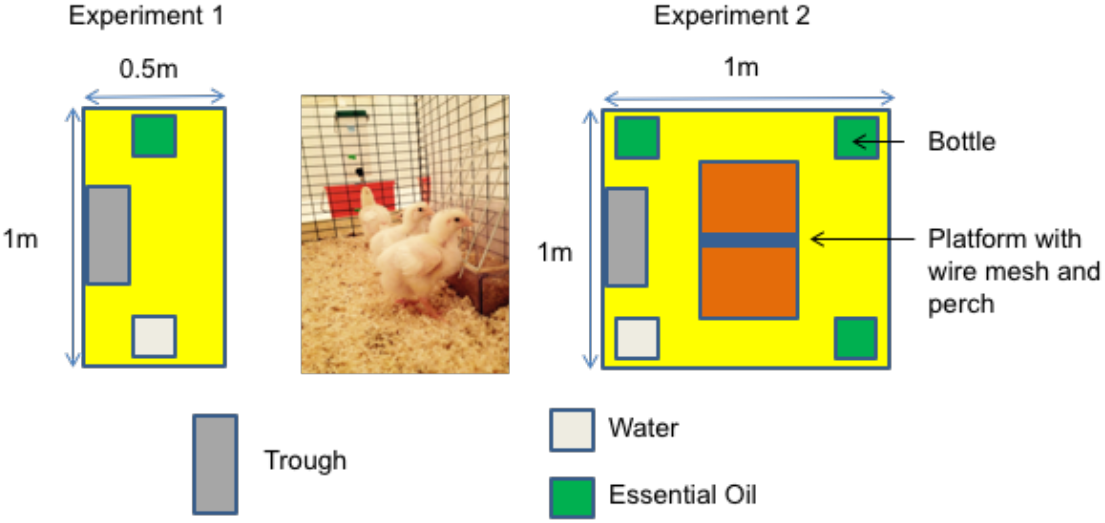
Experimental design

#### 2. Experiment 2

After incubator opening, 384 broiler chicks were either placed in pens (1m x 1m, 16 chicks/pen) in the C group (n = 192) or in the D group after 24h of the negative experience (n = 192). Before placement in pens, half of the chicks (n = 192) were randomly chosen to be macroscopically examined in order to define their quality scores as proposed by Tona *et al* (Tona et al., 2003). Only criteria focused on the retracted yolk (non-retracted: 0, retracted:12 (23% of total score)), navel area (not closed and discolored:0; not completely closed and not discolored: 6; completely closed and clean: 12 (23% of total score)), remaining membrane (very large membrane:0; large membrane: 4; small membrane: 8; no membrane: 12 (23% of total score)), and remaining yolk around the navel area (very large yolk: 0; large yolk: 8; small yolk: 12; no yolk: 16 (31% of total score)) were considered to establish a total score reported of 100%. Chicks were allocated to the W-C or W-D, or EO-C or EO-D group (six pens/group, 16 chicks/pen). Besides feed and water supplies, the three EO were provided together in EO groups (EO-C or EO-D) in free *ad libitum* access in the pens from D1 until D12 post-hatching. The three EO were placed together in the pens for the EO groups. Four bottles, one with water and three with each EO, were placed per pen (Fig. 1). The other half of the chicks (six pens of C group and six pens of D group) only had access to water in four bottles (W-C and W-D groups). As in the first experiment, the bottle position was changed every day for a week and every 2-3 days for the second week. At D13, the chickens were transferred in another livestock building and reared in standard conditions without EO access. The intake of water and of each EO was recorded at D1, D2, D4, D6, D9 and D12. Water and EO were changed each time of intake recording and supplemented if necessary between two measures of intake. The EO intake was expressed as the percentage of EO intake to the total liquid intake.

### Performance measurement

Body weight was measured at D0, D6, D9 and D13 in experiment 1, and at D0, D1, D6, D12, D19, D27 and D33 in experiment 2. Feed consumption was measured in each pen for the periods between D1-D6, D6-D9 and D9-D13 in experiment 1, between D0-D6, D6-D12, D12-D19, D19-D27 and D27-D33 in experiment 2, and used to calculate the feed conversion ratio (FCR). Twelve chickens per group (two/pen) were necropsied at D1 and D13 to measure the weight of the yolk sac and at D34 to measure the weight of the *Pectoralis major* muscle (experiment 2).

### Welfare status assessment (Experiment 2)

Several tests were used to measure fearfulness since this reactivity is supposed to be increased when birds experience stressful situations and this is why some of them are included in the Welfare Quality^®^ protocol (WelfareQuality, 2009).

#### Tonic immobility test

Tonic immobility is a behavioral response modulated by frightening situations and its duration is considered to be a measurement of the level of fearfulness (Forkman et al., 2007). Tonic immobility was induced by restraining the animal on its back: the longer the time needed for the bird to right itself (referred to as TI), the more fearful the bird was. Four 7-day-old chicks per pen were placed on their back in a U-shaped cradle and restrained for 10 sec and the duration of tonic immobility was recorded. If a chick failed to right itself after five min, a maximum score of 300 sec was recorded. If tonic immobility was not induced after five attempts, a score of 0 sec was recorded.

#### Novel object test

A novel object test was used to assess bird reactions to novelty with a protocol adapted from the Welfare Quality^®^ protocol. The novel object used was a 50–cm long and 3-cm wide stick with coloured bands. Five min after entering the pen, the observer placed the novel object on the litter between the trough and the bottles. The observer moved back 1.5 meter, kept standing, and counted every 30 sec for a total of two min the number of chicks at a distance of less than one chick length from the object and the number of chicks that pecked the object. This sampling was performed in each pen at 22 days of age.

#### Avoidance test

The avoidance distance test described in the Welfare Quality^®^ protocol was adapted to our experimental room for the assessment of the human-animal relationship. The observer entered the pen and kept standing close to the door since the pens were too small to allow walking without major disturbances to the chickens. The duration that was needed for three chickens at least (n = 12/pen) to come close to the observer (less than one meter) was recorded. This test lasted two min and was performed in each pen at 23 days of age.

### Health status assessment (Experiment 2)

General health status and the possible presence of respiratory, digestive, and musculoskeletal disorders were recorded each time that body weight was recorded. There was no evidence of hock burn nor of foot pad dermatitis, so only lameness was measured on D29 using the Welfare Quality^®^ gait scoring method. Gait scoring was performed by experts in four chickens per pen using score from 0 (normal gait) to 4 (severe abnormality, only able to walk few steps).

Immune system activity was assessed by measuring the antibody titers specific to the infectious bronchitis (IB) vaccine that were present in the serum of the chicks at hatching and at D13 and D34 after vaccination (Experiment 2). Antibody titers were determined by ELISA using the ID Screen^^®^^ IBV Indirect kit and the protocol described by the supplier (ID.vet, Grabels, France).

### Statistical analysis

Analyses were carried out using XLSTAT software (version 2015, Addinsoft, Paris, France). The effects of the delayed placement and EO supply on total liquid intake, water intake, total EO intakes, body and muscle weight and the FCR ratio were analyzed by ANOVA after having checked the normality of data distribution and the homogeneity of variances. When there are interactions between variables, the Fisher (LSD) test was used to determine the significant differences between groups. Because the data were not normally distributed and variances were not homogenous between groups, data on each EO intake, behavioral tests and gait scores were analyzed with non-parametric tests: the Kruskal-Wallis test for the group effect and the Mann-Whitney test for the comparison between the D and C groups for each period. The effects of periods on EO intake were analyzed with the non-parametric Friedman test. The Dunn test with the Bonferonni correction was used as a *post-hoc* test after Kruskall-Wallis and Friedman analyses. The clinical data and quality score of the chicks were analyzed by a Chi-squared test. Differences were considered to be significant when p-values were below 0.05 and not significant (NS) when p-values were above 0.1. The values are presented as means ± standard deviations or medians, quartiles, maximum, and minimum values.

## Results

### Spontaneous intake of one EO by chicks after a negative postnatal experience (Experiment 1)

The chicks drank significantly fewer liquids (water and EO) in the D group than in the C group whatever the group having access to water only or both water and EO but there was not EO effect within D or C groups (Table 2). They drank less water in the EO groups than in W groups and within these groups, they drank less water in the D groups than in the C group (Table 2). To overcome the effect of the liquid intake difference between D and C groups, the intake of EO was then normalized by reporting the intake of EO to the intake of liquids during each period analyzed.

**Table 2.**
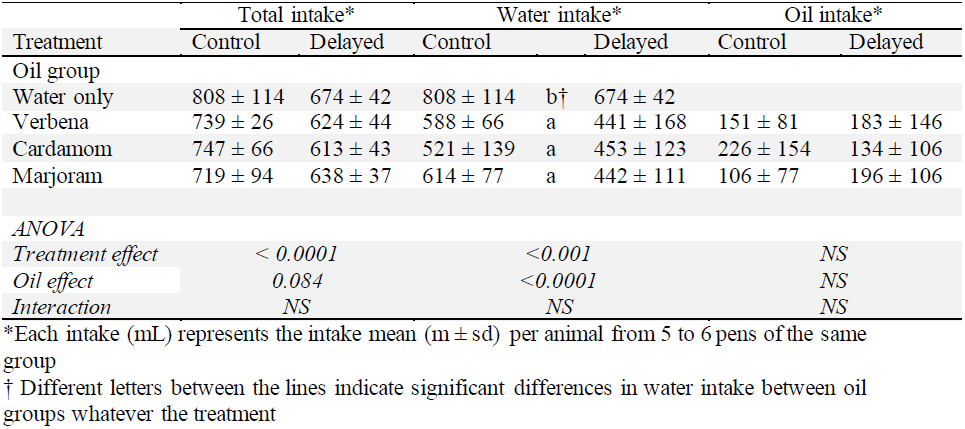
Liquid intakes in each group according to the treatment (Experiment 1)

There was a high variation in each EO intake between pens and for each group of chicks (C or D groups). However, there was a significant progressive increase in the EO intake over time for some EO (Figure 2). The intake of verbena EO was significantly higher for the period D9 to D13 compared to D1-D2 (Figure 2A). There were no significant differences in the intake of verbena EO for the D group during the period D1 to D13. For the C group, the intake of cardamom EO was higher for the period D6 to D13 compared to the intake for the period D2-D4 (Figure 2B). For the D group, the intake of cardamom EO was only higher for the period D9 to D13 compared to D2-D4. There were no significant differences over time in the intake of marjoram EO for both the C and D groups (Figure 2C).

**Figure 2.**
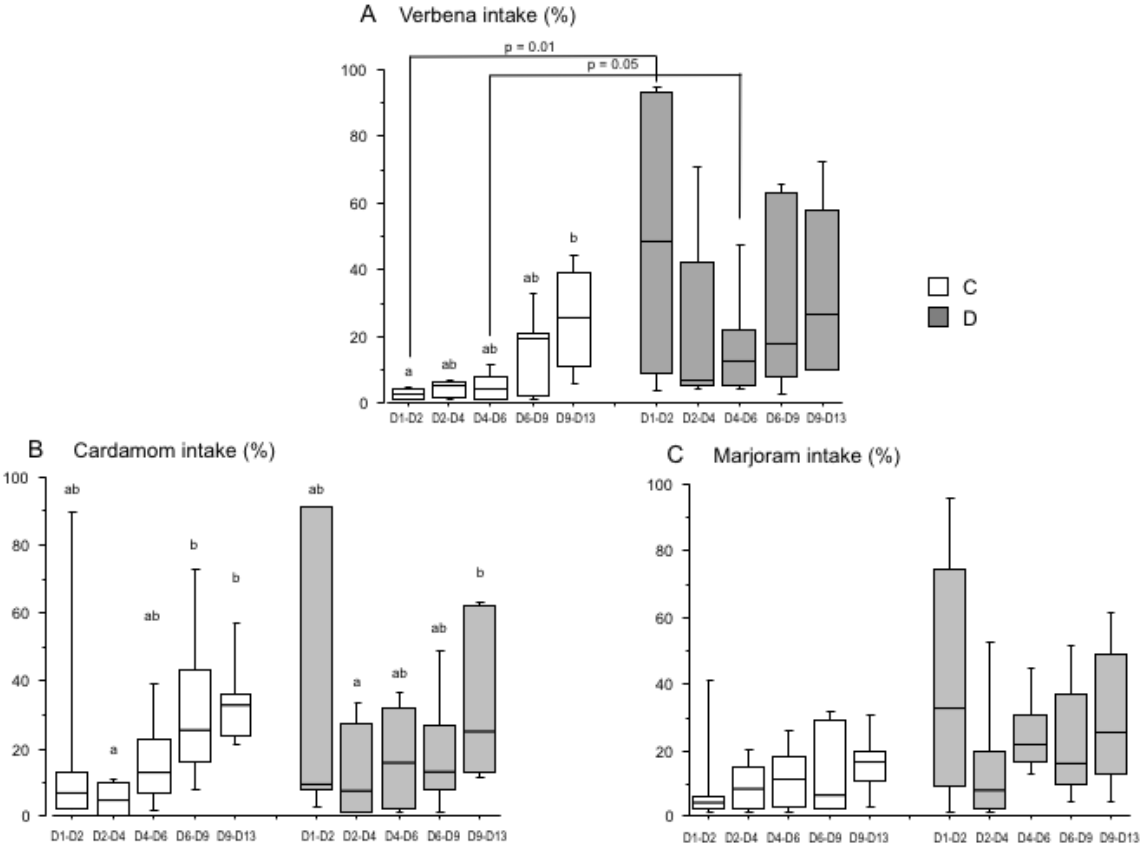
Essential oil intake by chicks over time (Experiment 1) Chicks were either directly placed in pens (C group; white) or delayed for 24h (D group; grey). The histograms show the box-plots and whiskers of EO intake (EO/(water + EO), %) for each group, verbena (A), cardamom (B), and marjoram (C). Different letters indicate significant differences in EO intake between periods of measurement for each group of chicks (Dunn test). The p-values indicate significant differences between the C and D groups within each period (Mann-Whitney test).

EO intake differed according to their postnatal treatment. The intake of verbena EO was significantly higher in the D group than in the C group for the periods D1-D2 and D4-D6, the amount of EO consumed was the highest between D1 and D2 (p = 0.01) (Figure 2A). The intake of cardamom EO was not significantly different between the D and C groups (Figure 2B). There was only a tendency for the D group to drink more marjoram EO than in the C group between D1 and D6 (25.4 ± 14.4 in the D group *vs* 10.9 ± 7.1 in the C group, p = 0.1) (Figure 2C). These results showed that the intake of verbena EO by chicks increased progressively over time according to their postnatal experience. The chicks spontaneously and rapidly drank more verbena EO when their placement in the rearing facility was delayed than when they were directly placed after hatching.

### Choice and spontaneous intake between three EO by chicks after a negative postnatal experience (Experiment 2)

In this experiment, the chicks had the choice to drink either water or any of the three EO used in the first experiment in addition to the water in free access in their pen. As in the first experiment, the chicks consumed significantly fewer liquids (water and EO) in the D group than in the C group for 12 days after hatching. They also drank less water in the EO groups than in the W groups independent of the postnatal treatment during this period (Table 3).

**Table 3.**
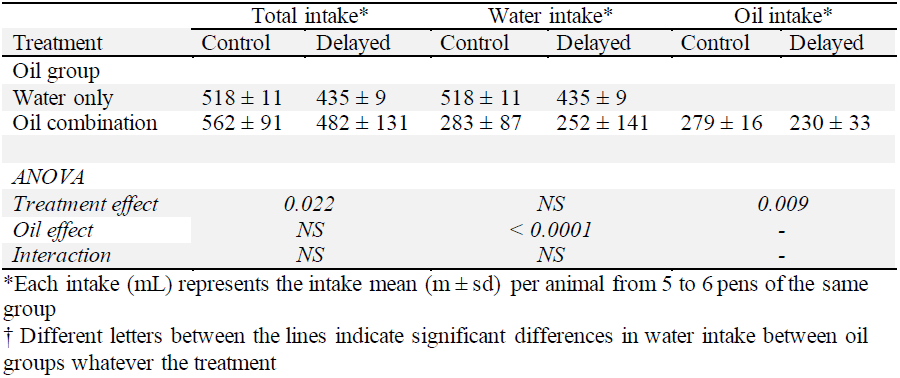
Liquid intakes in each group according to the treatment (Experiment 2)

As in the first experiment, there was a large variation in each EO intake between pens, and for both the C and D groups, there was a significant progression in the intake of EO between D1 and D12 (Figure 3A). There were no significant differences in the intake of verbena EO by the C group over time, but the intake by the D group increased significantly from the period D1-D2 to D6-D9 (Figure 3B). The intake of cardamom EO by the chicks increased progressively and significantly between D1-D2 and D6-D9 for the C group and from D2-D4 to D6-D9 for the D group (Figure 3C). It was the same for the intake of marjoram EO from the period D1-D2 to D4-D6 for both groups of chicks (Figure 3D).

**Figure 3.**
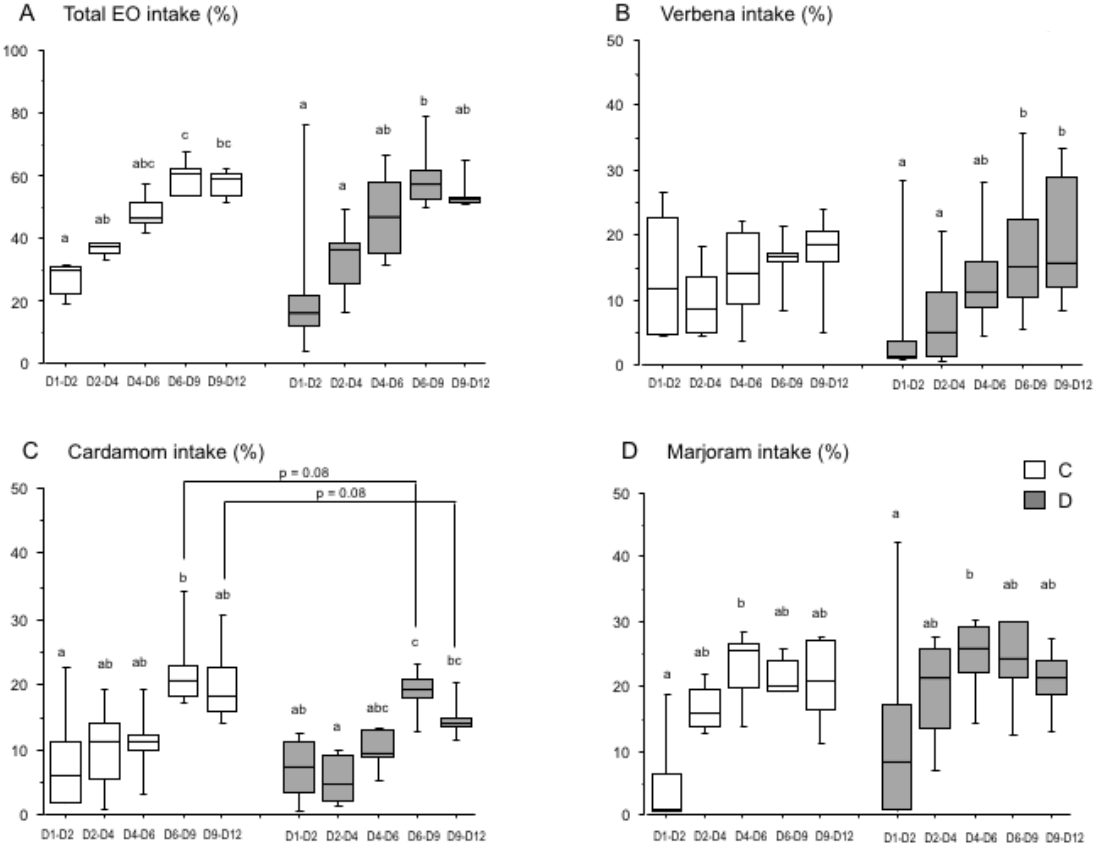
Essential oil intake by chicks over time (Experiment 2) Chicks were either directly placed in pens (C group, white) or delayed for 24h (D group; grey). The histograms show the box-plots and whiskers of EO intake (EO/(water + EO), %) for each group, the three EO (A), verbena (B), cardamom (C) and marjoram (D). Different letters indicate significant differences between periods for each group of chicks (Dunn test). The pvalues indicate significant differences between the C and D groups within each period (Mann-Whitney test).

The spontaneous intake of EO was different between the three EO available in pens depending on the postnatal treatment. The intake of cardamom EO was significantly lower in the D group than in the C group from D6 to D12 after hatching (16.4 ± 3.0 in the D group *vs* 21.1 ± 6.2 in the C group, p = 0.05), but only a tendency (p = 0.08) when comparisons were done between D6-D9 and D9-D12 (Figure 3C). However, the intake of verbena and marjoram EO was not significant between the C and D groups whatever the period of intake (Figures 3B and 3D). These results showed that when three EO were available simultaneously, the intake of EO by the chicks changed over time within each group of postnatal treatment, but EO were not differently chosen by chicks between groups except for the delayed chicks which drank less cardamom EO than control chicks.

### Effects of EO intake on chick performance

In experiment 1, the delay period of 24h before the placement of the chicks in the D group significantly reduced the chicks’ growth when they arrived in the building and until D13 (weight decrease of 14.8% less in the D group compared to the C group, p < 0.0001). The reduction in growth in the D group was not mitigated by EO intake (Figure 4A), but the FCR was significantly lower during the period of D6 to D9 in the EO groups than in the W groups, independent of the postnatal treatment (Figure 4B).

**Figure 4.**
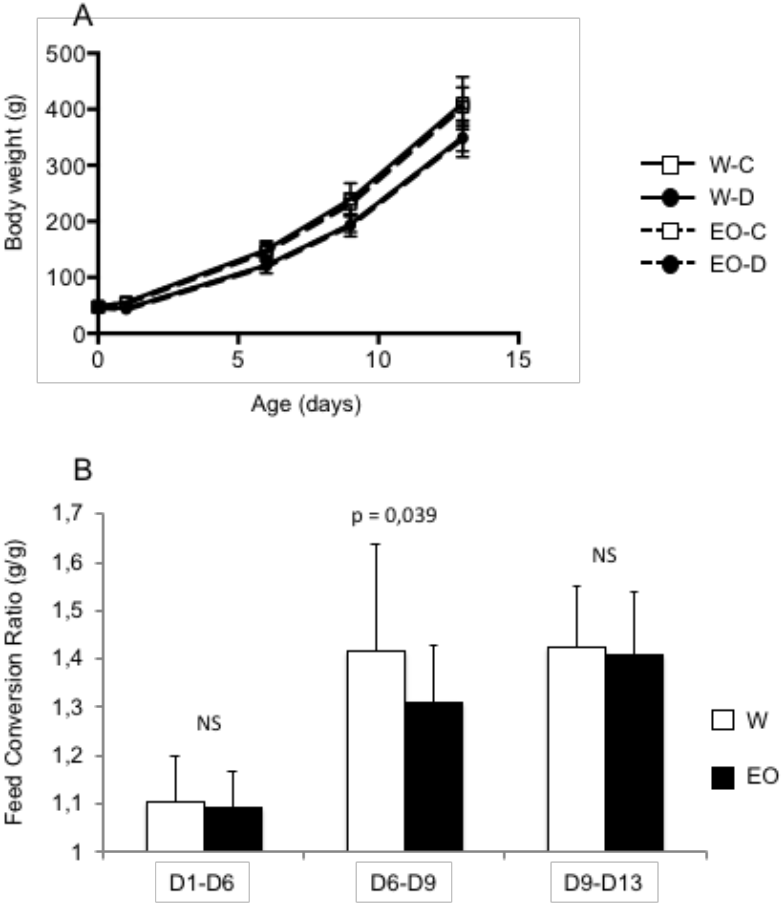
Chicken performance in experiment 1 Chicks were either directly placed in pens (C group) or delayed for 24h (D group), and had *ad libitum* access to only water (W) or to water and EO (EO). The curves show body weight for each group (A). The histograms show Feed conversion ratio (FCR) per period for chickens that had *ad libitum* access to only water (W) or to water and EO (EO) (B). The results express the mean and standard deviation. NS or p-values indicate statistical significance between the W and EO groups.

In experiment 2, 162 chicks among the 192 examined at T0, before any treatment and placement, had a global quality score superior to 36 points (out of 52 points, 70%) (class 1) and only 30 chicks had an inferior score (class 2). Chick weight at different times after hatching was not different in class 1 compared to class 2.

The size and the presence of vitellus in the chicks were not affected by the postnatal experience and EO ingestion at D13 and D34. Chicks in the D group had a very significant growth reduction from D1 to D34 (6.5%, p < 0,0001) (Figure 5A). The FCR in the D group was significantly impaired after an environment change (building and feed changes between D12-D19) compared to the C group, but it was the opposite during the following period D19-27 and there was no difference anymore after D27 (Figure 6). EO intake had no significant effect on chick growth and on FCR regardless of the postnatal experience. However, at D34, the *P. major* muscle yield was significantly increased in the chickens that had access to EO (Figure 5B), suggesting that EO intake has a positive effect on the growth rate of *P. major*.

**Figure 5.**
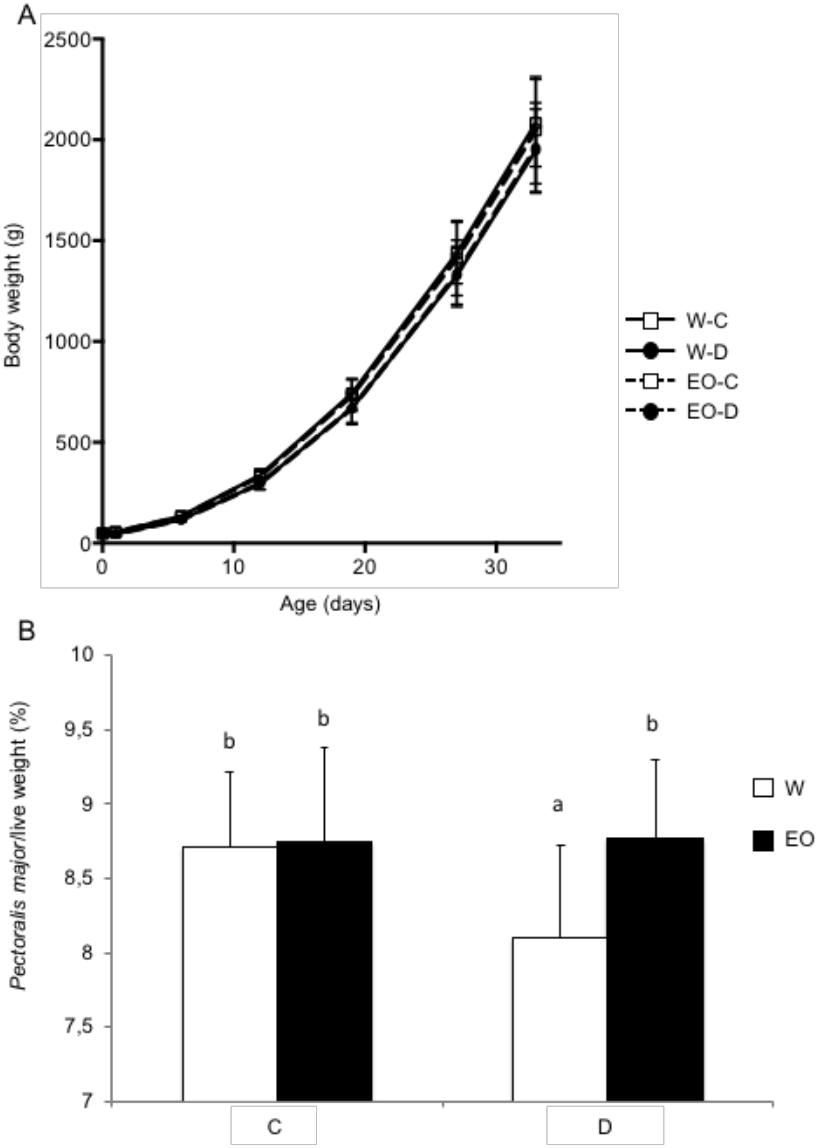
Chicken performance in experiment 2 Chicks were either directly placed in pens (C group) or delayed for 24h (D group), and had *ad libitum* access to only water (W) or to water and EO (EO). The curves show body weight for each group (A). The histograms show *Pectoralis major* weight at 34 days of age for chickens that had *ad libitum* access to only water (W) or to water and EO (EO) (B). The results express the mean and standard deviation. NS or p-values indicate statistical significance between the W and EO groups.

**Figure 6.**
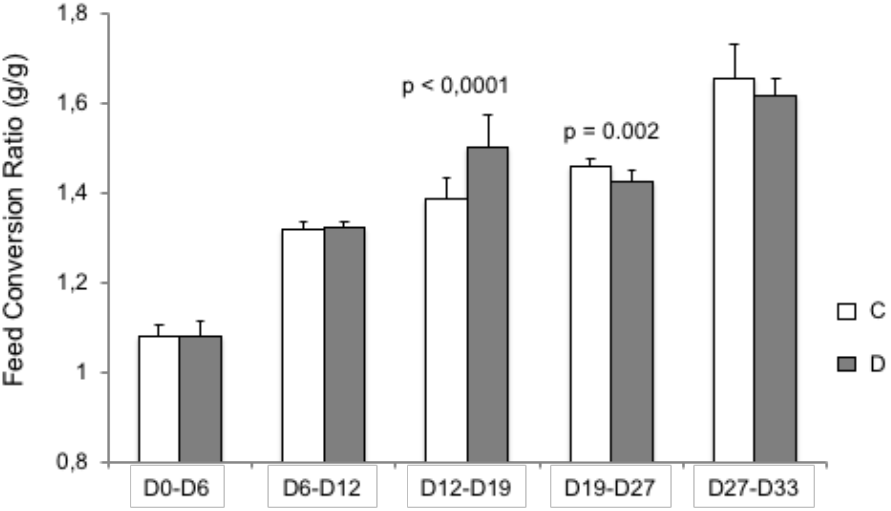
Chicken performance in experiment 2 The histograms show Feed conversion ratio (FCR) per period for chicks either directly placed in pens (C group) or delayed for 24h (D group), and that had *ad libitum* access to only water or to water and EO. The results express the mean and standard deviation. NS or p-values indicate statistical significance between C and D groups.

### Effects of EO intake on chick welfare

Delayed placement and EO supply had no effect on tonic immobility duration (54.2 s ± 42.3) or on the number of attempts needed to induce this behavior (1.6 ± 0.8).

During the novel object test, the mean number of chickens close to the object or pecking at it in each group was not influenced by the delayed placement or by EO supply regardless of the scan period (0 sec, 30 sec, 60 sec, 90 sec, 120 sec). The average number of chickens close to the object was 5.2 ±1.6 and the average number of chickens pecking at it was 1.2 ±2.0.

The mean number of chickens close to the observer in each group was not influenced by the delayed placement or by EO supply over most the scan period (0 sec, 30 sec, 60 sec, 90 sec, 120 sec). This number was lower in the EO groups 60 sec after test starting (4.6 ± 1.6 in EO pens *vs* 6.5 ± 2.0 in W pens, p = 0.03) and the average number over all scan periods tended to be lower in the EO pens compared to the W pens (4.5 ± 1.4 in EO pens *vs* 5.9 ± 1.6 in W pens, p = 0.06).

### Effects of EO intake on chick health (Experiment 2)

The prevalence of respiratory and digestive disorders was not different between the C and D groups, or between chicks that were in quality score class l at hatching (good quality) compared to class 2, or between chicks with access or not to EO. The gait score (2.21 ± 0.59) was affected neither by the negative postnatal experience nor by EO supply. The global mortality rate in the experiment was 2.6%. It was associated with a lower chick quality score at hatching (37.3 ± 6.9 in dead animals vs 45.1 ± 6.8, p = 0.04 in others). Chicks died either of heart attack (n = 3) or were euthanized because of severe locomotor disorders (n = 5).

Regarding the reactivity of the immune system, the antibody response after vaccination against IB was analyzed. The antibody response dropped after hatching and was not different between the C and D groups or between chicks with access or not to EO (Figure 7).

**Figure 7.**
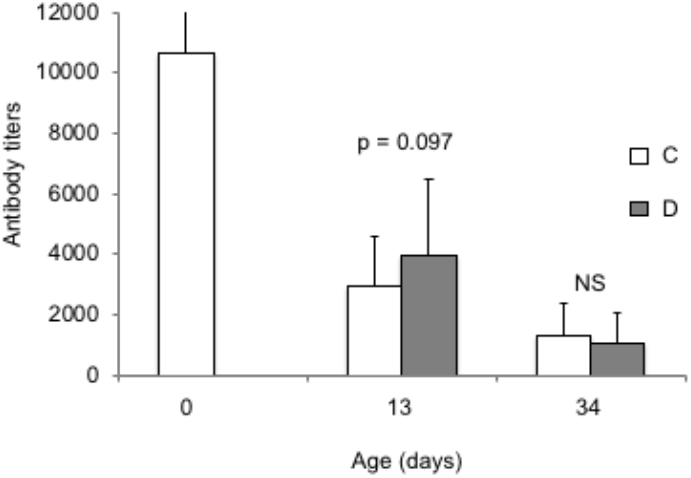
Antibody titers after infectious bronchitis vaccination of chicken (Experiment 2) The histograms show antibody titers in serum from chicks either directly placed in pens (C group) or delayed for 24h (D group), and that had *ad libitum* access to only water or to water and EO. At day 0, the antibody titer was analyzed at hatching time before the delayed treatment. The results express the mean and standard deviation. NS or p-values indicate statistical significance between C and D groups.

## Discussion

The present study investigated the capacity of chicks to select EO and to consume EO after their exposure to a negative postnatal experience related to the delay between their hatching and their transportation to the rearing facilities. The chicks consumed significantly fewer liquids (water and EO) in the delayed group than in the control group for 12 days after hatching, possibly in relationship with the significant reduction in body weight induced by the delayed placement for the D group. Regarding the EO intake itself, a considerable variation was found in EO intake between the chick pens of the delayed group, particularly in the days following the stressful event (D1-D2) and it was also true for the control group to a lesser extent. A progressive increase in EO intake was observed over time for most of the EO. In the first experiment, when the chicks had the choice to consume water or one EO, the cardamom EO intake increased significantly over time and it started earlier for the C group (D6) than for the D group (D9). It was also the case for the verbena EO intake in the C group (D9) but it was not the case for the D group or the marjoram EO for both groups. Many animals can use medication by selecting and eating specific plants (de Roode et al., 2013a). The process involved in medication behavior is complex and the involvement of innate versus learned behavior is discussed whether it concerns therapeutic or prophylactic medication (de Roode et al., 2013b;Moore et al., 2013). This discussion is usually restricted to immune defenses against parasites and the process of learning about food containing secondary plant compounds (de Roode et al., 2013b;Moore et al., 2013). In our study, we chose to introduce EO in water and in supplement to water available to satisfy the chickens’ thirst. So the spontaneous intake of EO by chicks was independent of thirst. The immediate intake of verbena EO by the D group could suggest an innate behavior of medication whereas the progressive intake of this EO by the C group and of cardamom EO for both groups could suggest a learning process over time.

In the second experiment, when the three EO were simultaneously available, there was also a progressive increase in EO intake over time for both groups, except for the verbena EO intake in the C group. Actually, it was the opposite result of the first experiment, the verbena EO intake by the D group increased progressively over time whereas the intake was immediate and constant for the C group. We can assume that it was more difficult for the chicks in the D group to learn from post-ingestion signals since these signals were probably merged because of the simultaneous provision of the three EO. It has been shown from diets with different energy levels that chicks are able to build preferences when they have acquired experiences of the post-ingestion cues of the diets (Bouvarel et al., 2008). This suggests that chick choices between several EO would need a previous experience with each EO separately. However, we do not have any explanation about the immediate and persistent intake of verbena EO by the C group in experiment 2.

When the EO were presented separately (experiment 1), the chicks spontaneously consumed verbena EO over the period of EO access from D1 to D13 and in significantly higher amount in the D group than in the C group from D1 to D6. There was a tendency for the D group to consume more marjoram EO from D1 to D6 than the C group. These results showed that chicks were able to make the spontaneous choice of drinking verbena EO, and possibly marjoram EO, immediately after their negative postnatal experience and for a week. During that period, the control group drank a small amount of these EO.

Several conditions have been identified to define the behavior of SM: (1) infection or discomfort induces SM behavior, (2) SM improves the fitness of infected animals, and (3) SM behavior is costly to non-infected animals (Singer et al., 2009;de Roode et al., 2013a). In chickens, one study reported the preference of lame chickens for a feed supplemented with an anti-inflammatory and analgesic drug (Carprofen) rather than the same feed without the drug (Danbury et al., 2000). This study suggested that lame broilers found a benefit in eating feed supplemented with carprofen and may select carprofen for its analgesic properties. The control chickens tended to avoid feed supplemented with carprofen, suggesting an aversion to this drug. In our study, the high intake of verbena EO by the delayed chicks and its low intake by the control chicks during the 6 days after the negative postnatal experience suggest that delayed chickens may select verbena EO for its properties. The antioxidant, anti-inflammatory, sedative, and digestive effects of lemon verbena are well reported in *in vitro* and *in vivo* studies and more recently the beneficial effect of this EO on muscle damage after exhaustive exercise was described (Buchwald-Werner et al., 2018). The exposure of the chicks to the combination of feed and water deprivation, temperature changes, and unpredictable shaking may explain their choice to consume verbena EO. Likewise, the tendency of the D group to select marjoram EO may be related to its antioxidant and hepatoprotective properties (Bina and Rahimi, 2017), which could help the chicks to overcome their delayed placement.

In contrast, the delayed chicks drank less cardamom EO after 6 days compared to the control chicks when the EO was in combination with the two other EO. Yet, in addition to antioxidant and anti-inflammatory activities, cardamom EO has antispasmodic and gastroprotective activities (al-Zuhair et al., 1996;Jamal et al., 2006). The beneficial effect on performance of a diet supplemented with cardamom EO has been reported in broilers, and a positive effect on the blood cholesterol profile was shown (Omidi et al., 2015). In our study, the lower consumption of cardamom EO in the D group than in the C group could suggest that the costs/benefits of cardamom EO intake for the D group was too high. This behavior has been reported in monarch butterfly fitness costs after using antiparasitic plant chemicals (Tao et al., 2016) and in ruminants (Villalba et al., 2017). A model developed by Choisy and de Roode (Choisy and de Roode, 2014) suggests that animals evolve phenotypic plasticity when parasite risk is low to moderately high and genetically fixed medication when parasite risk becomes very high. Although many animals use secondary chemicals to recover health, medication behaviors can result in substantial fitness costs, which are associated with the concentration and composition of biologically active secondary metabolites (Tao et al., 2016). In our study, we estimate the amount of EO ingested by the chicks to be from 6 to 12 µg/g/chick between D1 and D12. This is very low compared to the amount of EO ingested when they are integrated in the diet, about 100 mg/kg of feed, which corresponds to around 10 mg of EO/chicken per day at 12 days of age (Adaszynska-Skwirzynska and Szczerbinska, 2017).

The SM behavior should improve the fitness of infected or uncomfortable animals. In our study, the postnatal experience of the combination of feed and water deprivation, temperature changes, and unpredictable shaking of the transportation boxes before the placement in rearing houses had a significant and long lasting effect on the chickens’ growth until the age of slaughter (Day 34). It had a temporary negative effect on FCR when an unexpected event occurred such as the transfer of chicken in another building. This is in agreement with previous studies focused on one type of postnatal experience (Bigot et al., 2003;Gonzales et al., 2003;Bergoug et al., 2013;Zhao et al., 2014;Jacobs et al., 2016;de Jong et al., 2017), but it can differ in chickens according to the age, as well as the type and the duration of the stressors. For example, food restriction during the first week of a chick’s life had beneficial effects on performances and resistance to infection disease (Zulkifli et al., 1994). In our study, the delayed placement did not have significant long-lasting effects on chicken welfare and health, maybe because health disorders were limited to leg problems and not related to any infectious diseases. However, the altered FCR observed when an unexcepted event occurred in delayed chickens suggests that they were less efficient for their performance than for maintaining their welfare and health in our experimental conditions. The EO intake did not have any significant effect on growth, but had a positive effect on the *P. major* muscle yield. Positive effects on chicken performance were reported elsewhere using EO, however differing from the ones used in our study, in drinking water at similar concentrations (Adaszynska-Skwirzynska and Szczerbinska, 2017;2018).

In conclusion, our study showed that chicks could select EO according to their postnatal experience. The selection and the intake of EO varied with the chicks’ age, which suggests that adding a mix of EO in a determined concentration into the diet or into the water supply would not allow to chicks to adapt their intake to their needs. It would be more appropriate to give chickens access to a diversity of feed and non-nutritive extracts with medicinal properties throughout their life. These results were obtained in broiler chicks whose genotype has been selected for their high growth rate. Although domestication is thought to increase stress tolerance, the genetic selection of broiler chickens has been detrimental to their adaptive immunity and subsequently their resistance to pathogens. The present results are in favor of a conserved SM behavior which could allow the chickens to individually balance their performance, health, and welfare. Encouraging studies on SM could contribute to more sustainable rearing practices and veterinary medicines.

## Acknowledgments

We would like to thank the staff of the Avian Experimental Unit (INRA, UE 1295 Pôle d’Expérimentation Avicole de Tours, 37380 Nouzilly, France) for producing the chickens and for their assistance in experiments, S. Crochet, E. Cailleau-Audouin and E. Baéza from the Avian Biology and Poultry Science Unit (UMR BOA, INRA, Université de Tours, 37380 Nouzilly, France) and P. Constantin (UMR PRC, INRA, CNRS, IFCE, Université de Tours, 37380 Nouzilly, France) for their help in experiments, and G. Mauboussin, O. Honcharova, A. Yildirim, the students involved in the study. We also thank M. Derval and C. Ingraham for their expertise in aromatherapy and self-medication practices with livestock animals. The authors are grateful to L. Bignon and I. Bouvarel (Institut Technique de l’Aviculture, F-37380 Nouzilly, France) for helpful discussions and S. Beauclercq (UMR BOA, INRA, Université de Tours, 37380 Nouzilly, France) for revision. This research was supported by a grant from the Integrated Management of Animal Health Metaprogram of INRA for the « GISA-WHELP » project (www.gisa.inra.fr/en). The English was reviewed by M. Pinier (Carpe Sensum, translation and revision language).

## Author contributions

LG, CL and AC conceived and designed the experiments. LG and CL supervised the study and wrote the first draft of the manuscript. All the authors contributed to the technical work, the data analyses and to manuscript revision, read and approved the submitted version.

## Data availability statements

The raw data supporting the conclusions of this manuscript will be made available by the authors, without undue reservation, to any qualified researcher.

